# Microdiversity and phylogeographic diversification of bacterioplankton in pelagic freshwater systems revealed through long-read amplicon sequencing

**DOI:** 10.1101/2020.06.03.133140

**Authors:** Yusuke Okazaki, Shohei Fujinaga, Michaela M. Salcher, Cristiana Callieri, Atsushi Tanaka, Ayato Kohzu, Hideo Oyagi, Hideyuki Tamaki, Shin-ichi Nakano

## Abstract

Freshwater ecosystems are inhabited by members of cosmopolitan bacterioplankton lineages despite the disconnected nature of these habitats. The lineages are delineated based on >97% 16S rRNA gene sequence similarity, but their intra-lineage microdiversity and phylogeography, which are key to understanding the eco-evolutional processes behind their ubiquity, remain unresolved. Here, we applied long-read amplicon sequencing targeting nearly full-length 16S rRNA genes and the adjacent ribosomal internal transcribed spacer sequences to reveal the intra-lineage diversities of pelagic bacterioplankton assemblages in 11 deep freshwater lakes in Japan and Europe. Our single nucleotide-resolved analysis, which was validated using shotgun metagenomic sequencing, uncovered 7–101 amplicon sequence variants for each of the 11 predominant bacterial lineages and demonstrated sympatric, allopatric, and temporal microdiversities that could not be resolved through conventional approaches. Clusters of samples with similar intra-lineage population compositions were identified, which consistently supported genetic isolation between Japan and Europe. At a regional scale (up to hundreds of kilometers), dispersal between lakes was unlikely to be a limiting factor, and environmental factors were potential determinants of population composition. The extent of microdiversification varied among lineages, suggesting that highly diversified lineages (e.g., Iluma-A2 and acI-A1) achieve their ubiquity by containing a consortium of genotypes specific to each habitat, while less diversified lineages (e.g., CL500-11) may be ubiquitous due to a small number of widespread genotypes. The lowest extent of intra-lineage diversification was observed among the dominant hypolimnion-specific lineage (CL500-11), suggesting that their dispersal among lakes is not limited despite the hypolimnion being a more isolated habitat than the epilimnion. Our novel approach complemented the limited resolution of short-read amplicon sequencing and limited sensitivity of the metagenome assembly-based approach, and highlighted the complex ecological processes underlying the ubiquity of freshwater bacterioplankton lineages.

## Introduction

Microbial phylogeography is the study of the diversification and distribution of microorganisms across space and time, and offers insights into eco-evolutionary processes that generate and maintain the ubiquity and diversity of microbial populations. However, our understanding of microbial phylogeography is far behind that of macroorganisms [1] and has not yet achieved a general consensus, as evidenced by the fact that the old tenet of microbiology “Everything is everywhere, but the environment selects” remains a matter of debate [2]. The next key step is accurately profiling the diversity of environmental microbial assemblages, which is challenging because they are dominated by organisms that are recalcitrant to cultivation [3, 4]. The rapid advances of sequencing and bioinformatics technologies have provided novel opportunities for cultivation-independent, high-resolution analysis that may effectively address this long-standing topic.

For prokaryotes, the current *de facto* standard approach for phylogenetic profiling of an environmental community is high-throughput amplicon sequencing of the 16S rRNA gene. However, in exchange for its universality, the phylogenetic resolution of the 16S rRNA gene is limited, allowing organisms to be resolved to the genus, but not species level [5, 6]. Furthermore, the phylogenetic resolution is usually limited by the short reads (150–300 bp paired-end) generated by Illumina sequencers, which is the most commonly used sequencing platform and can sequence only a portion of the 16S rRNA gene (∼1500 bp). To achieve finer phylogenetic resolution, the more variable ribosomal internal transcribed spacer (ITS) region, located between the 16S and 23S rRNA genes, is a common alternative marker. The ITS sequence can differentiate ecologically distinct intra-lineage populations that cannot be resolved using the 16S rRNA gene. ITS-based microdiversities across space, time, and environmental gradients have been reported for members of marine *Pelagibacter* [7, 8] picocyanobacteria [9–11] and freshwater *Limnohabitans* [12–14], *Polynucleobacter* [15], and *Synechococcus* [16]. In contrast to the 16S rRNA gene, ITS lacks a universal primer and a comprehensive database, and is too variable for determining diversity across lineages; for example, the length of ITS varies by hundreds of base pairs within a phylum [17]. Therefore, the ITS is ideally sequenced along with the adjacent 16S rRNA gene to achieve finer phylogenetic resolution and broad taxonomic classification [18].

The technical limitations of short-read platforms have recently been tested using long-read sequencing platforms, namely Pacific Biosciences (PacBio) and Oxford Nanopore. These platforms can sequence amplicons of the complete 16S rRNA gene [19–22] and even the entire rRNA gene operon, including the ITS and small and large subunit rRNA genes (>4000 bp), of prokaryotes [23–25] and microbial eukaryotes [26–28]. A major drawback of long-read sequencing is its higher per-base error rate compared to short-read sequencing. With the PacBio platform, this issue can be solved through construction of a circular consensus sequence (CCS), in which individual amplicon molecules are sequenced many times using circularized library templates that provide consensus-sequence error correction [20]. When combined with a quality-filtering process based on per-base quality scores, the error rate of a CCS-generated amplicon can be reduced to a level comparable to those of short-read platforms [19, 24, 27] allowing analysis at single-nucleotide resolution [29].

Genomic average nucleotide identity (ANI) is another promising approach for high-resolution phylogenetic profiling of prokaryotes, with ANI>95% considered a robust threshold for species-level classification [30–32]. ANI is often applied to a metagenome-assembled genome (MAG), a draft-quality genome reconstructed from shotgun metagenomic sequences, providing the opportunity to perform cultivation-independent, genome-resolved phylogenetic analysis. However, the MAG-based approach suffers from several technical limitations: First, the number of samples is practically limited due to the high cost of metagenomic sequencing. Second, the analysis requires reconstruction of a high-quality MAG, which is challenging for bacterial lineages with low abundance (i.e., low sequencing coverage) or those harboring highly microdiversified genotypes [33, 34]. Third, a MAG often lacks a 16S rRNA gene due to the difficulty of reconstructing such a highly conserved region [35], making it difficult to link the MAG with 16S rRNA gene-based phylogenies. Long-read amplicon sequencing can avoid these limitations and provide a complementary approach to MAG-based analysis that can achieve high-resolution phylogenetic profiling of an environmental microbial assemblage.

Lakes are physically disconnected ecosystems, and thus offer a good model for investigating the diversification and phylogeographic processes of microbes. Whereas 16S rRNA gene-based phylogenies have been used to characterize the cosmopolitan bacterioplankton lineages that are ubiquitously dominant in freshwater systems [36–38], analyses at finer phylogenetic resolution have been used to identify intra-lineage microdiversity and phylogeography. A global-scale investigation of microdiversity was conducted for the PnecC subcluster of the genus *Polynucleobacter* (Betaproteobacteria), a cosmopolitan freshwater bacterial group, using the ITS and *glnA* gene as markers, which showed distance–decay patterns and global-scale niche separation of subgroups adapted to different thermal conditions [15]. Genome-resolved studies focusing on other ubiquitous lineages, such as LD12 (Alphaproteobacteria) and acI (Actinobacteria), have also shown that geographically isolated lakes are generally inhabited by distinct species (i.e., ANI<95%) [39, 40] Meanwhile, genomes sharing ANI>95% were isolated from lakes located up to hundreds of kilometers apart and separated by the Alps [41, 42] and even between lakes on different continents [40, 43, 44]. No global phylogeographic patterns at the phylogenetic resolutions of ITS [45] and ANI [46] were observed for *Microcystis*, ubiquitous bloom-forming freshwater cyanobacteria. Due to the sparseness of data at fine phylogenetic resolution across habitats and lineages, the following questions remain open: (i) How much microdiversity exists for each bacterioplankton lineage within and among lakes? (ii) Does microdiversity exhibit any phylogeographic patterns following distance–decay relationships or environmental gradients? (iii) Are phylogeographic patterns similar or distinct among microbial lineages?

To address these questions, the present study first applied single nucleotide-resolved long-read amplicon sequencing targeting the nearly full-length 16S rRNA gene and ITS regions (∼2000 bp) to investigate the microdiversity and phylogeographic patterns of multiple freshwater bacterioplankton lineages among multiple lakes. Sampling was performed at pelagic sites of deep (>70 m) oligo-mesotrophic freshwater lakes. Compared with shallow freshwater habitats, deep lakes are fewer in number, but are characterized by larger water volume, longer water retention time, and older age. Thus, the pelagic microbial communities of deep lakes are expected to be less strongly influenced by disturbances from terrestrial and sedimental inputs and to show more robust spatial and temporal distributions, allowing analysis of limited samples to better represent their diversification, dispersal, and historical processes. We explored nine Japanese and two European lakes and performed analyses at both the regional and intercontinental scales. In each lake, samples were collected from two water layers, the surface mixed layer (epilimnion) and the oxygenated hypolimnion (water layer below the thermocline). The oxygenated hypolimnion is generally found in deep oligo-mesotrophic holomictic lakes and is inhabited by specific bacterioplankton lineages [38]. While inhabitants of the epilimnion may migrate over long distances using neighboring shallow lakes and ponds as “stepping stones” [45, 47], hypolimnion inhabitants are more likely to be isolated due to the limited occurrence of their habitat. Therefore, we hypothesized that hypolimnion-specific lineages would be more deeply diversified among lakes than epilimnion-specific lineages.

## Methods

### Sample collection

Samples were collected from nine Japanese and two European perialpine lakes (Fig. 1 and Table 1). Physicochemical details of the lakes are provided in Table S1. In each lake, two water layers, the epilimnion and oxygenated hypolimnion, were sampled at a pelagic station during the stratification period (Table 1). Samples of the perialpine lakes were collected in October 2017 using 0.22-μm pore-size polyethersulfone filter cartridges (Millipore, Sterivex SVGP01050) following prefiltration through a 5.0-μm pore-size polycarbonate filter (Whatman, cat. no. 111113). At least 2 L of lake water was filtered for each sample and the filter cartridge was stored at −20°C until further processing. The filter was manually removed from the Sterivex cartridge and processed using the PowerBiofilm DNA Isolation Kit (MoBio Laboratories) for extraction of DNA. Samples from the Japanese lakes were collected during a previous study in 2015 for short-read 16S rRNA gene amplicon sequencing [38], and the remaining DNA extracts were used for the present study. In addition, DNA samples collected in October 2010 from Lake Biwa [48] were used as temporal replicates for comparison of samples taken in 2010 and 2015 from the same lake.

**Table 1.** Main characteristics of lakes sampled in the present study. Detailed data are available in Table S1.

| Lake | Region | Origin | Surface area (km<sup>2</sup>) | Surface elevation (m) | Maximum depth (m) | Water retention time (y) | Mixing regime | Trophic state | Sampling date | Sampling depth (m) |
| --- | --- | --- | --- | --- | --- | --- | --- | --- | --- | --- |
| Mashu (MA) | Japan (Hokkaido) | Caldera | 19.2 | 351 | 211 | 200 | Dimictic | Oligotrophic | 2015.8.26–29 | 1.5, 200 |
| Kusharo (KU) | Japan (Hokkaido) | Caldera | 79.6 | 121 | 118 | 12 | Dimictic | Oligotrophic | 2015.8.25 | 1.5, 50 |
| Toya (TO) | Japan (Hokkaido) | Caldera | 70.7 | 84 | 180 | 9 | Dimictic | Mesotrophic | 2015.9.24 | 5, 150 |
| Inawashiro (IN) | Japan (Honshu) | Tectonic | 103.3 | 514 | 94 | 5.4 | Monomictic | Oligotrophic | 2015.10.30 | 5, 80 |
| Chuzenji (CH) | Japan (Honshu) | Dammed | 11.8 | 1,269 | 163 | 5.9 | Dimictic | Oligotrophic | 2015.10.21 | 5, 120 |
| Sai (SA) | Japan (Honshu) | Dammed | 2.1 | 900 | 72 | 1.6 | Monomictic | Oligotrophic | 2015.9.8 | 10, 50 |
| Motosu (MO) | Japan (Honshu) | Dammed | 4.7 | 900 | 122 | 6.5 | Monomictic | Oligotrophic | 2015.9.7 | 10, 100 |
| Biwa (BI) | Japan (Honshu) | Tectonic | 670.3 | 85 | 104 | 5.5 | Monomictic | Mesotrophic | 2015.12.9, (2010.10.19)† | 2, 60 (5, 50)† |
| Ikeda (IK) | Japan (Kyushu) | Caldera | 10.9 | 66 | 233 | 33.5 | Holo-oligomictic | Mesotrophic | 2015.9.17 | 5, 70 |
| Zurich (ZU) | Europe (Switzerland) | Glacial | 88.0 | 406 | 136 | 1.4 | Holo-oligomictic | Mesotrophic | 2017.10.25 | 5, 60 |
| Maggiore (MG) | Europe (Italy/Switzerland) | Glacial | 213.0 | 193 | 370 | 4.1 | Holo-oligomictic | Oligo-mesotrophic | 2017.10.16 | 5, 100 |

† Sampling date and depths in 2010 are shown in parentheses.

**Figure 1.**
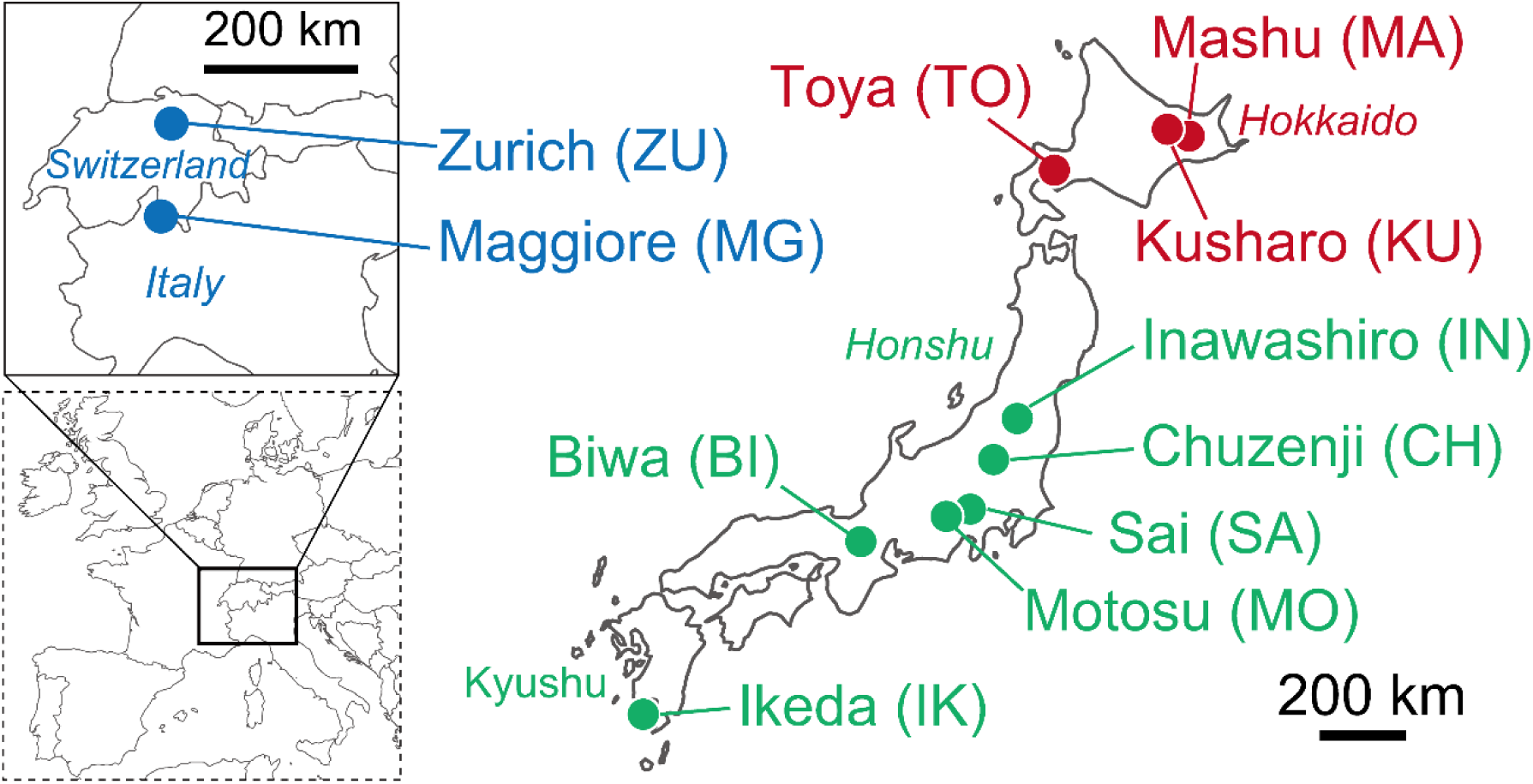
Locations of the lakes sampled in the present study. Colors indicate regions: green, Honshu and Kyushu islands; red, Hokkaido island; blue, Europe.

### Polymerase chain reaction (PCR), library preparation, and sequencing

To amplify the nearly full-length 16S rRNA gene and ITS sequences, we used the primer set 27Fmod [5′-AGRGTTTGATYMTGGCTCAG-3′][49] and 23Sr-mod [5′-RGTTBYCYCATTCRG-3′]. The latter primer was based on the widely used 23Sr (also known as 23S-125r) primer [5′-GGGTTBCCCCATTCRG-3′], which targets the ITS side of the 23S rRNA gene [12, 18, 50–52], and was modified to cover broader taxonomic groups (Table S2). For multiplexing, a 16-base index sequence was prepared for each sample and conjugated to the 5′ end of both the forward and reverse primers. PCR was performed in 25-μL reactions containing 2 μL of template DNA (1–5 ng μL^-1^) using the Blend Taq Plus (TOYOBO, Osaka, Japan) buffer system. Cycling conditions were as follows: initial denaturation at 94°C for 180 s, followed by 35 cycles of amplification (denaturation at 94°C for 45 s, annealing at 50°C for 45 s, extension at 72°C for 90 s) and a final extension at 72°C for 180 s. Amplification of approximately 2000-bp amplicons was confirmed through agarose gel electrophoresis. For each sample, at least two reaction mixtures were prepared and pooled after amplification to obtain sufficient DNA and mitigate potential PCR biases. The resulting PCR products were purified using AMPure XP beads (Beckman Coulter), quantified with the Qubit dsDNA HS Assay kit (Thermo Fisher Scientific), and pooled equimolarly. The sequencing library was prepared using the SMRTBell Template Prep Kit 1.0 following the Procedure and Checklist 2 kb Template Preparation and Sequencing protocol (Pacific Biosciences, Inc.; document version, PN001-143-835-08). Sequencing was performed on the PacBio RSII sequencer using P6-C4 chemistry and five SMRT (single-molecule real-time) cells with a 4-hour video length. Demultiplexed CCSs were generated using the RS_ReadsOfInsert.2.1 protocol in SMRT Analysis software version 2.3.0 with the following settings: Minimum Full Passes=2, Minimum Predicted Accuracy=90, and Minimum Barcode Score=22 in Symmetric Barcode Mode. The raw CCS reads were deposited under accession number PRJDB9651.

### Analysis of sequencing outputs

The 159,262 CCS reads (average length=1,951 bp) obtained from the 24 samples were analyzed using the DADA2 v. 1.12.1 package [53] with R v. 3.4.4 software (http://www.R-project.org/), which is a pipeline for assigning reads to single nucleotide-resolved amplicon sequence variants (ASVs). We followed the workflow previously designed for CCS-based long-read amplicon sequencing [29]: The demultiplexed FASTQ files of CCS reads were processed by sequentially applying the removePrimers, filterAndTrim, derepFastq, learnErrors, dada, makeSequenceTable, and removeBimeraDenovo functions (the R script used here is available in the Supplementary Text). The resulting table mapped 28,153 reads from 24 samples to 742 ASVs (Supplementary Dataset).

The 742 ASV sequences thus obtained (including both 16S rRNA gene and ITS) were aligned against the SILVA SSU 132 Ref NR database [54] using the SINA Aligner v. 1.2.11 webserver [55] with default parameters. Based on the alignment results, the 16S rRNA gene region was extracted from each ASV and clustered using the --cluster_fast command in VSEARCH v. 2.8.0 software [56]. Clusters generated using 97% and 100% identity thresholds were designated as operational taxonomic unit (OTU) and SSU-ASV, respectively. A representative sequence for each OTU was determined using the --consout option. Each OTU was taxonomically classified using the SINA Aligner v. 1.2.11 webserver with reference to the SILVA SSU 132 Ref NR database. In addition, each OTU was annotated based on the nomenclature for freshwater bacterioplankton proposed by Newton et al. [36], which was applied using the ‘--search’ option in the SINA v. 1.2.11 stand-alone tool against the original ARB [57] database provided by Newton et al. [36]. Finally, ASVs were assigned back to OTUs and SSU-ASVs using the --usearch_global command in VSEARCH software with 97% and 100% identity thresholds, respectively (Supplementary Dataset). For unknown reasons, the number of reads assigned to each sample varied considerably: from 190 (epilimnion of Lake Zurich) to 4515 (hypolimnion of Lake Biwa), with a median of 1114.5 (Supplementary Dataset). To make the best use of the available data, we used all reads for analysis and did not perform rarefaction. For each OTU, samples with <20 assigned reads were not included in subsequent analysis to minimize biases caused by a low number of reads. To avoid biases introduced by uneven sequencing depths among the samples, our investigation focused on the dominant ASVs and did not address the overall richness or rare ASVs.

For each OTU, the pairwise Bray–Curtis dissimilarities of ASV composition were calculated among samples using the vegan v. 2.5-2 package [58] of the R software. Hierarchical clustering of the data was performed using the hclust function with the ‘methods=“ward.D2”’ option, and the results were visualized using the ComplexHeatmap v. 2.2.0 package [59] of R software.

### Assessment of methodological performance through metagenomic read mapping

To evaluate the performance of our single nucleotide-resolved analysis, we tested whether the single nucleotide polymorphisms (SNPs) predicted from ASVs can be reproduced in shotgun metagenomic reads. We focused on sequences of the CL500-11 lineage in Lake Biwa, and used the assembly and read mapping results generated from a sample collected in the hypolimnion of the lake in September 2016 [60]. Reads mapped to the contig containing the rRNA gene operon of CL500-11 were analyzed using IGV v. 2.4.10 software [61] to determine the base frequency for each nucleotide position.

## Results and Discussion

### Phylogenetic composition of long-read amplicons

Overall, our analysis generated 742 ASVs assigned to 441 SSU-ASVs and 155 OTUs (Supplementary Dataset). The results revealed the diversity of sequence-discrete populations within lineages sharing >97% or even 100% 16S rRNA gene sequence identity, which could not have been observed using single nucleotide-resolved short-read amplicon sequencing analysis of the same samples [38]. The sequences were affiliated with ten phyla, four of which accounted for >95% of total reads. Samples from the epilimnia were dominated by the phyla Actinobacteria (68.3% of reads), Proteobacteria (18.3%), and Bacteroidetes (10.8%), while the phyla Actinobacteria (50.8%), Chloroflexi (34.5%), and Proteobacteria (9.3%) dominated the hypolimnion samples (Supplementary Dataset). For further analysis, we selected 11 OTUs (hereafter “dominant lineages”) that were ubiquitous (detected in more than eight samples) and abundant (>300 reads; 1.1% of the total). The dominant lineages collectively accounted for 85.4% of total read abundance in the study, and included one OTU in Chloroflexi (CL500-11), one in Alphaproteobacteria (LD12), and nine in Actinobacteria (acI-A1, acI-A6, acI-A7, acI-B1, Iluma-A1, Iluma-A2, acIV-B, and two acI-C2) (Supplementary Dataset).

### Intra-lineage microdiversity and phylogeographic patterns among lakes

Each of the dominant lineages harbored 5–34 SSU-ASVs and 7–101 ASVs (Supplementary Dataset). Despite the large number of ASVs detected in each lineage, the majority of ASVs were rare, with only a few ASVs accounting for most reads in each sample. The average proportion of the most abundant ASV in a sample ranged from 43.8% (acI-A1) to 84.9% (acI-C2-e), and that of the three most abundant ASVs ranged from 78.5% (acI-B1) to 100% (acI-C2-e) (Fig. 2). Based on the ASV composition of each sample (Fig. S1), Bray–Curtis dissimilarity matrices among samples were generated for each lineage (Fig. S2), and the overall pattern across lineages was evaluated by averaging the matrices of the 11 dominant lineages (Fig. 3). The results indicated that five clusters shared closely related ASV compositions (Fig. 3); while lakes in Europe and Hokkaido formed separate clusters, samples from other Japanese lakes clustered by water layer (i.e., epilimnion and hypolimnion), except for samples from Lake Inawashiro, which clustered independently (Fig. 3). Other exceptions were the hypolimnion of Lake Chuzenji (included in the Hokkaido cluster), and the epilimnion of Lake Motosu (clustered separately from all other samples) (Fig. 3).

**Figure 2.**
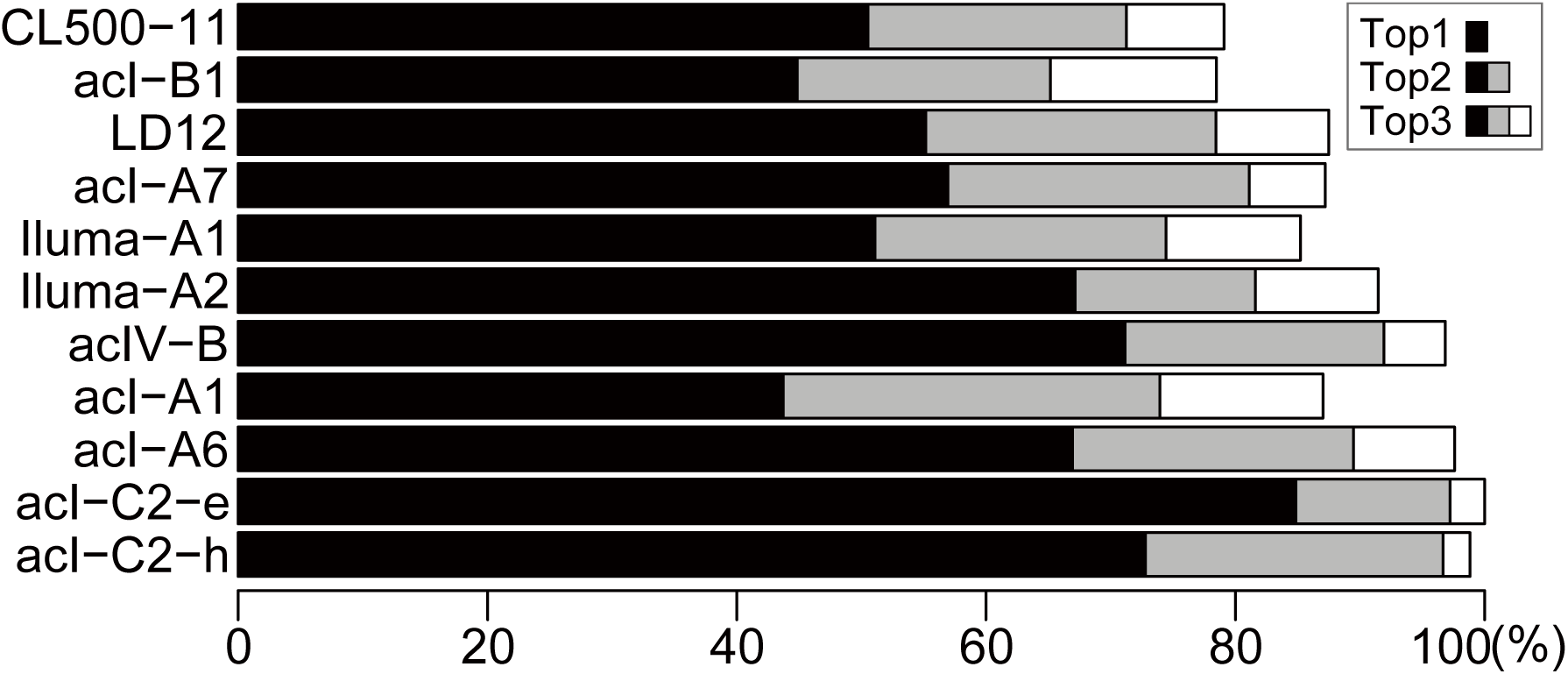
Proportion of reads representing the three most abundant amplicon sequence variants (ASVs), shown as the average value among all samples. Black bars indicate the average read percentage of the most abundant ASV for each lineage. The average percentages of the second and third most abundant ASVs are indicated with stacked gray and white bars. Note that acI-C2 comprised two different operational taxonomic units that were specific to the epilimnion (acI-C2-e) and hypolimnion (acI-C2-h), respectively.

**Figure 3.**
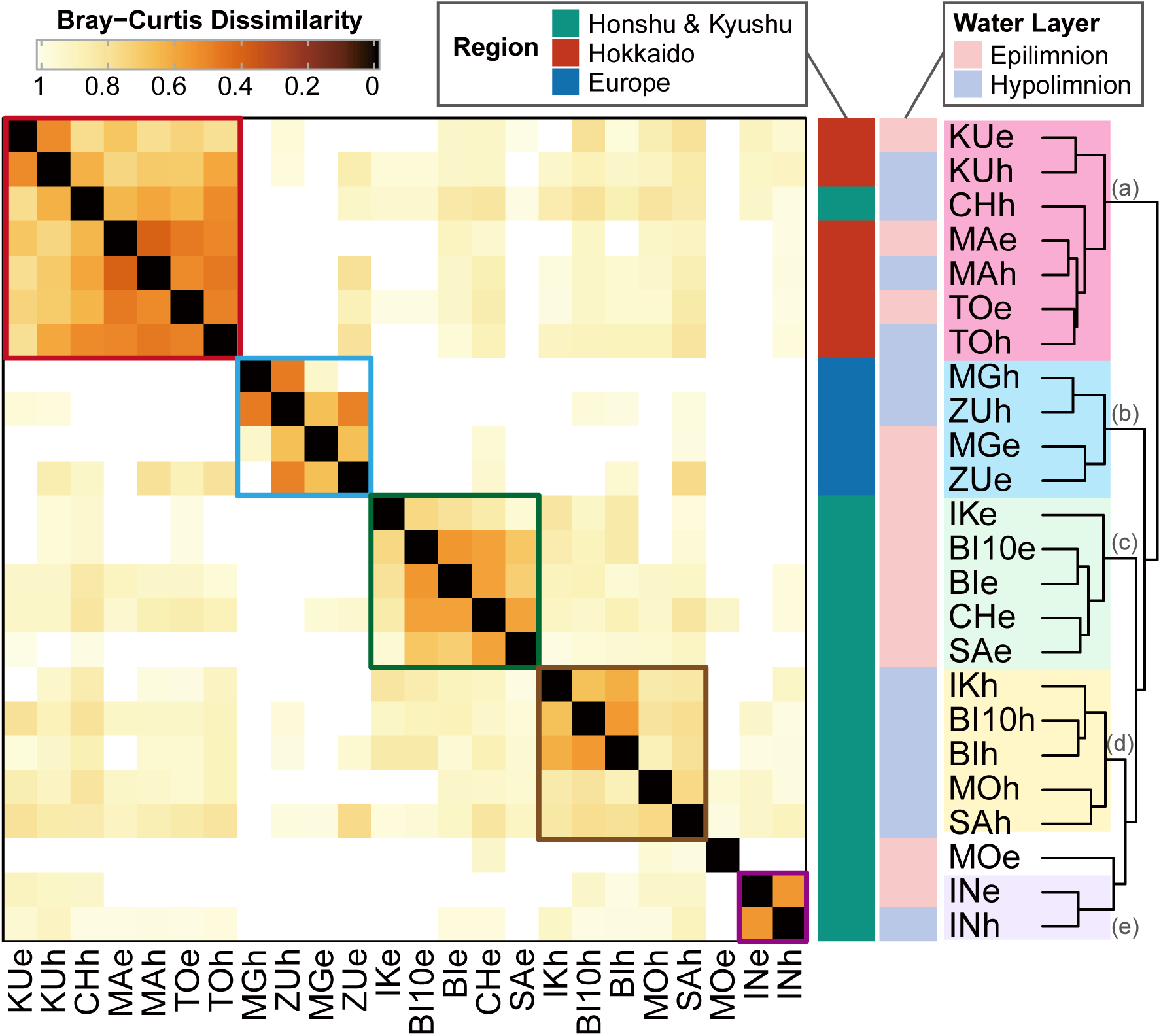
Clustering of samples based on the Bray–Curtis dissimilarity of amplicon sequence variant composition generated by averaging the values for the 11 most dominant lineages. The matrices for individual lineages are provided in Fig. S2. Sample names follow the abbreviations shown in Fig. 1, with a suffix indicating the water layer: e, epilimnion; h, hypolimnion. The temporal replicate collected in Lake Biwa in 2010 is abbreviated “BI10”. Five clusters were identified, grouping samples from (a) Hokkaido, (b) Europe, (c) Honshu and Kyushu epilimnia, (d) Honshu and Kyushu hypolimnia, and (e) Lake Inawashiro.

Clusters that were separated between Japan and Europe, as well as between Hokkaido and Honshu islands in Japan, were observed in most individual lineages (Fig. S2), indicating the universality of these trends. Planktonic organisms in inland waters can migrate between unconnected habitats on aerosols or due to animal and human activities [62]. Our results suggest that the migration of bacterioplankton across lakes is limited by distance. It should be noted that the observed phylogeographic pattern is a consequence not only of dispersal limitation, but also of successful colonization by migrating populations. The finding that few ASVs dominated lineages within each sample (Fig. 2) supports the hypothesis that members of the same lineage (i.e., those sharing >97% 16S rRNA gene similarity) have overlapping ecological niches; therefore, the environmental capacity for supporting multiple genotypes at the same time and place is limited. This hypothesis evokes the theory of priority effects, in which an increase in migrant genotypes due to genetic drift is limited by a high frequency of indigenous genotypes occupying the same niche [63]. Based on these results and assumptions, we propose that migration of lake bacterioplankton between Japan and Europe is unlikely, but that migration occurs at distances of up to hundreds of kilometers with a sufficiently high frequency to overcome priority effects.

The existence of epilimnion- and hypolimnion-specific clusters rather than distance–decay relationships among lakes in Honshu and Kyushu (Fig. 3) suggests that local environments are stronger factors in determining the population composition within the region than dispersal and random genetic drift. We speculate that the inclusion of the hypolimnion of Lake Chuzenji within the Hokkaido cluster (Fig. 3) might reflect environmental factors. Due to its high elevation, Lake Chuzenji is the coldest (with hypolimnetic temperature permanently at 4°C or lower) and the only dimictic lake in Honshu (Tables 1 and S1), and these conditions might have selected for cold-tolerant or psychrophilic populations that are also dominant in Hokkaido lakes. The unique microbial population in Lake Inawashiro (Fig. 3) may have resulted from environmental sorting in the past. While this lake is a typical oligotrophic monomictic lake today (Tables 1 and S1), it was previously acidic (pH<5.0) due to volcanic inflow and has experienced rapid neutralization (pH=6.6–7.0) over the last 30 years [64]. Because pH is one of the main drivers of lake microbial community composition [65], the present genotype diversity in the lake could have resulted from a bottleneck during the acidic period. Together, these results suggest that environmental sorting is a potential determinant of intra-lineage population composition in lake bacterioplankton, especially among habitats located within hundreds of kilometers.

Previous studies have investigated temporal trends in the intra-lineage population composition of freshwater bacterioplankton, demonstrating that the community consists of both persistent and transient populations [13, 66, 67]. In line with this finding, the temporal replicates collected in 2010 and 2015 in Lake Biwa revealed that the dominant ASVs were shared between the years, although some ASVs observed in 2010 were absent in 2015 and vice versa (Fig. S3). Remarkably, the temporal replicates from Lake Biwa were most closely related to each other based on the dissimilatory matrix (Fig. 3), indicating that the temporal change in the lake was less significant than the differences among lakes. Therefore, the observed phylogeographic pattern (Fig. 3) may be robust regardless of sampling time. Because our temporal analysis was limited to Lake Biwa, the results motivated further investigation of the potentially vast spectrum of bacterioplankton microdiversity across seasons and years in a lake, which will provide further insights into the mechanisms driving co-existence and turnover in multiple intra-lineage genotypes.

### Microdiversity and phylogeographic patterns of the most abundant hypolimnetic lineage (CL500-11)

In addition to the overall phylogeographic pattern, analysis of individual lineages provided a more detailed perspective on their microdiversification. Here, we focus on CL500-11 (‘*Ca*. Profundisolitarius’), a cosmopolitan bacterioplankton lineage that was dominant in the oxygenated hypolimnia of deep freshwater lakes [68, 69]. Although CL500-11 was almost exclusively detected in the hypolimnion, it was among the most abundant (6,084 reads; 21.6% of the total) and ubiquitous (detected in nine out of 11 lakes) lineages in the present study. We identified a total of 48 ASVs, among which four were dominant (ASV_1, 2, 5, and 10) and collectively accounted for 72.6% of CL500-11 reads in the Japanese lakes, and one (ASV_13) exclusively dominated European lakes and accounted for 99.2% of reads from those sites (Figs. 4 and S1). Between the ASVs observed in Japan and Europe, 5 bp mismatches were found in the 16S rRNA gene, along with at least 15 bp mismatches and a 1 bp gap in the ITS region (Fig. 4). These results further support our finding that bacterioplankton inhabiting European and Japanese lakes are genetically distinct.

**Figure 4.**
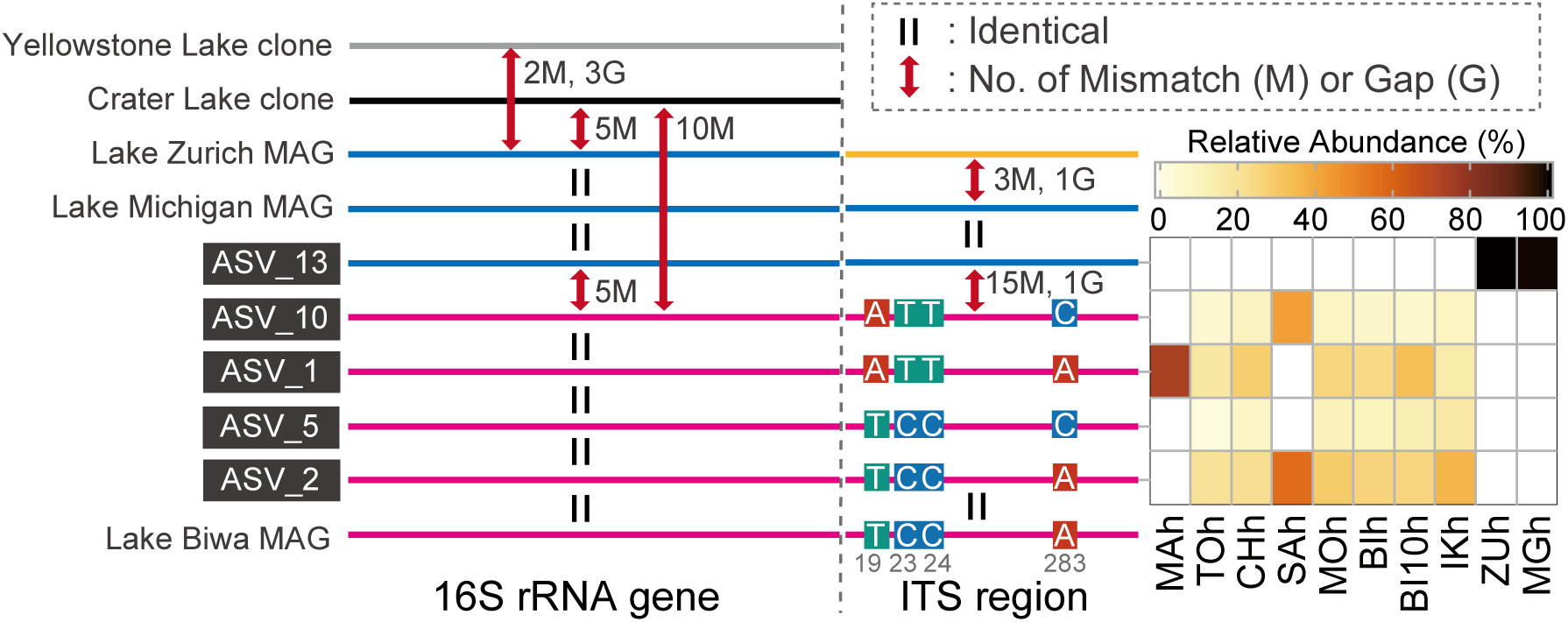
Schematic illustration of mismatches and gaps found in the 16S rRNA gene and internal transcribed spacer (ITS) sequences of the CL500-11 lineage, including the five dominant amplicon sequence variants (ASVs) and five publicly available sequences (see main text for details). Nucleotide variations among Japanese sequences (shown in pink) are displayed, with four variable base positions in the ITS sequence represented by gray numbers. Other mismatches and gaps are indicated with red arrows, and the same sequence type is indicated by the same color. The heat map indicates the relative abundances of ASVs within the CL500-11 lineage in each sample. Abbreviations for sample names follow those in Fig. 3.

The four predominant Japanese ASVs shared identical 16S rRNA gene sequences and differed at only four variable base positions in the ITS region (Fig. 4). Each of these ASVs was detected in at least five lakes (Fig. 4), resulting in a relatively uniform population composition across Japan compared with other dominant lineages (Figs. 5 and S2). Such a low degree of diversification was unexpected and contrary to our original hypothesis that hypolimnion-specific lineages would be more deeply diversified among lakes than epilimnion inhabitants due to their limited opportunities for dispersal. Instead, our results suggest that the preference for deep water is not a major factor limiting the diversification and dispersal processes of lake bacterioplankton. One may argue that their actual genomic diversity may be even lower, as different ASVs can be derived from multiple rRNA gene operons within the same genome. However, we ruled out this possibility based on the ASVs occurring independently; for example, only one of these ASVs (ASV_1) was detected in Lake Mashu (Fig. 4).

**Figure 5.**
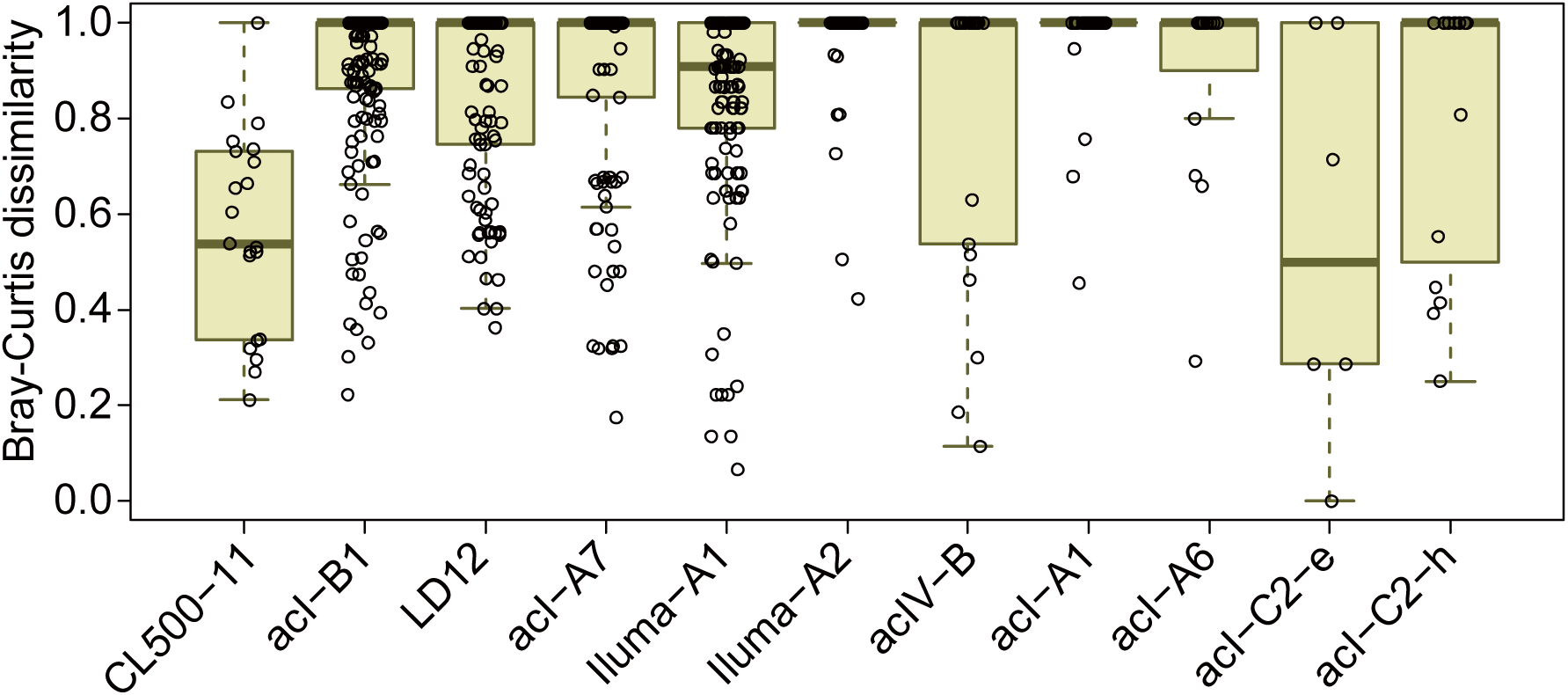
Distribution of pairwise Bray–Curtis dissimilarity for amplicon sequence variant (ASV) compositions among the nine Japanese lakes. Each point represents the dissimilarity between a pair of the samples, and their distribution is represented by a boxplot for each lineage. Note that dissimilarity is 1 when there are no shared ASVs between two samples, and thus many points are concentrated on the top of the boxes. The original pairwise dissimilarity matrices are available in Fig. S2. Note that acI-C2 comprised two different operational taxonomic units that were specific to the epilimnion (acI-C2-e) and hypolimnion (acI-C2-h), respectively.

A previous MAG-based analysis demonstrated that at least three species-level (ANI>95%) clusters exist within CL500-11: MAGs from Lake Biwa, Lake Michigan, and the Caspian Sea were distinct (ANI<95%) from each other, while those from Lake Zurich were found in both the Lake Michigan and Caspian Sea clusters [43]. The finding that MAGs from Europe and North America shared ANI>95% is in contrast to our finding that CL500-11 populations from different continents (Japan and Europe) were genetically distant from each other. Interestingly, the 16S rRNA gene and ITS sequence of the MAG from Lake Michigan [70] was identical to the European ASV in the present study (Fig. 4), further supporting their intercontinental genetic connectivity between Europe and North America. However, more information is required to draw a firm conclusion about their global-scale microdiversity and phylogeographic patterns, as evidence of unobserved sequence diversity has been found among the CL500-11 lineage. The ITS sequence of a CL500-11 MAG collected in Lake Zurich in May 2013 [43] was not identical to our European ASV (Fig. 4), despite the sequence of the MAG containing complete matches to the primers used in the present study. In North America, two full-length 16S rRNA gene sequences of CL500-11 were retrieved from Crater Lake (accession number=AF316759) [71] and Yellowstone Lake (HM446117) [72], and both exhibited 5 bp mismatches or gaps compared to the European ASV and to MAGs from Lake Michigan and Lake Zurich (Fig. 4). Moreover, three partial 16S rRNA gene sequences with at least one mismatch relative to our European ASV were detected in other European perialpine lakes (Lakes Annecy, Bourget, and Geneva) [73] (see Figure S10 in Okazaki et al. [38]). We should note that the number of reads assigned to CL500-11 was considerably lower in European lakes (379 reads) than in Japanese lakes (5705 reads) due to uneven sample number and sequencing throughput between the two regions (Supplementary Dataset). Thus, it is possible that rare sequence types of CL500-11 in European lakes could not be detected in our analysis due to insufficient data. Together, these findings indicate that the vast sequence space of the CL500-11 lineage remains underexplored both spatially and temporally.

### Comparison of diversification patterns among lineages

In contrast to the limited diversification of CL500-11 among Japanese lakes, Iluma-A2 and acI-A1 represented the opposite extreme, as their population compositions were unique for most lakes (Figs. 5, S1, and S2). Other dominant lineages exhibited intermediate degrees of diversification (Fig. 5). The wide degree of diversification among lineages may be due to differences in dispersal characteristics, such as the likelihood of aerosolization [74, 75]. Alternatively, the extent of diversification may reflect an ecological strategy of a lineage to exploit its genomic diversity. That is, lineages with relatively uniform population compositions among lakes (e.g., CL500-11) may harbor a few widespread genotypes that can dominate a broad range of habitats, whereas highly diversified lineages (e.g., Iluma-A2 and acI-A1) may be composed of many specialist genotypes that collectively colonize a wide range of environments. The latter case assumes partitioning of functional capabilities among genotypes within a lineage to minimize the amount of genetic material required per cell in a given niche [14, 41, 76]. Indeed, a high degree of microdiversification, mainly related to carbon metabolism, was reported among closely related acI-A1 strains with 100% identical 16S rRNA gene sequences, and even within the ANI>95% species boundary (e.g. ‘*Ca*. Planktophila versatilis’ and ‘*Ca*. Planktophila dulcis’) [39]. The microdiversity and ecological strategies underlying the ubiquity of freshwater bacterioplankton lineages are worth further exploration. The key step in such research is linking the 16S rRNA gene and ITS sequences with genomic and functional diversity, which requires assembly of high-quality genomes (i.e., those including the rRNA operon) that may be obtained through cultivation-based approaches [39, 41, 42] or long-read shotgun metagenome sequencing [77].

### Methodological consideration and future perspectives

Two major challenges are associated with the long-read sequencing platform: read accuracy and read throughput. In response to the former challenge, we targeted the 16S rRNA gene and ITS sequences but excluded the adjacent 23S rRNA gene, as a longer insert would result in (i) lower per-base quality of CCS due to fewer rounds of subreads and (ii) a smaller number of error-free reads at the same per-base error rate. To test the accuracy and sensitivity of our analysis, we focused on the CL500-11 lineage in the hypolimnion of Lake Biwa—the OTU and sample with the greatest number of reads (2606 reads) in the present study—and investigated whether the SNPs expected from ASVs could be reproduced through metagenomic read mapping. Although the metagenomic sample was collected a year after the main sampling period for the present study [60], metagenomic reads (i.e., randomly fragmented total DNA) reflect real sequence variance in the environment and thus are a good reference for assessing the performance of our approach. We identified 30 base positions with SNPs by aligning all 24 ASVs of CL500-11 detected in the sample (Fig. 6A). The results showed that among all observed SNPs, 78.9% were common to both ASV and mapping analyses, whereas only 5.3% were expected based on ASV but undetected through mapping (Fig. 6A and B). Such high reproducibility was remarkable because most SNPs expected based on ASVs were rare (occurring in <10% of the reads) (Fig. 6A). Further, the proportions of major SNPs were almost identical in ASV and mapping analyses (Fig. 6C). Notably, the original metagenomic study [60] assembled only one (ASV_2) of the ASVs present in the lake (Fig. 4). This limitation is due to the assembler generating a consensus high-quality contig to represent a lineage rather than fragmented assemblies reflecting different microvariants [33, 34, 78]. Overall, we demonstrated that, given a sufficient number of reads, our approach successfully recovered the variants and proportions of single nucleotide-resolved sequence diversity in the environment and is sensitive enough to detect minor sequence types with a low rate of false positives, which would be overlooked by the MAG-based approach.

**Figure 6.**
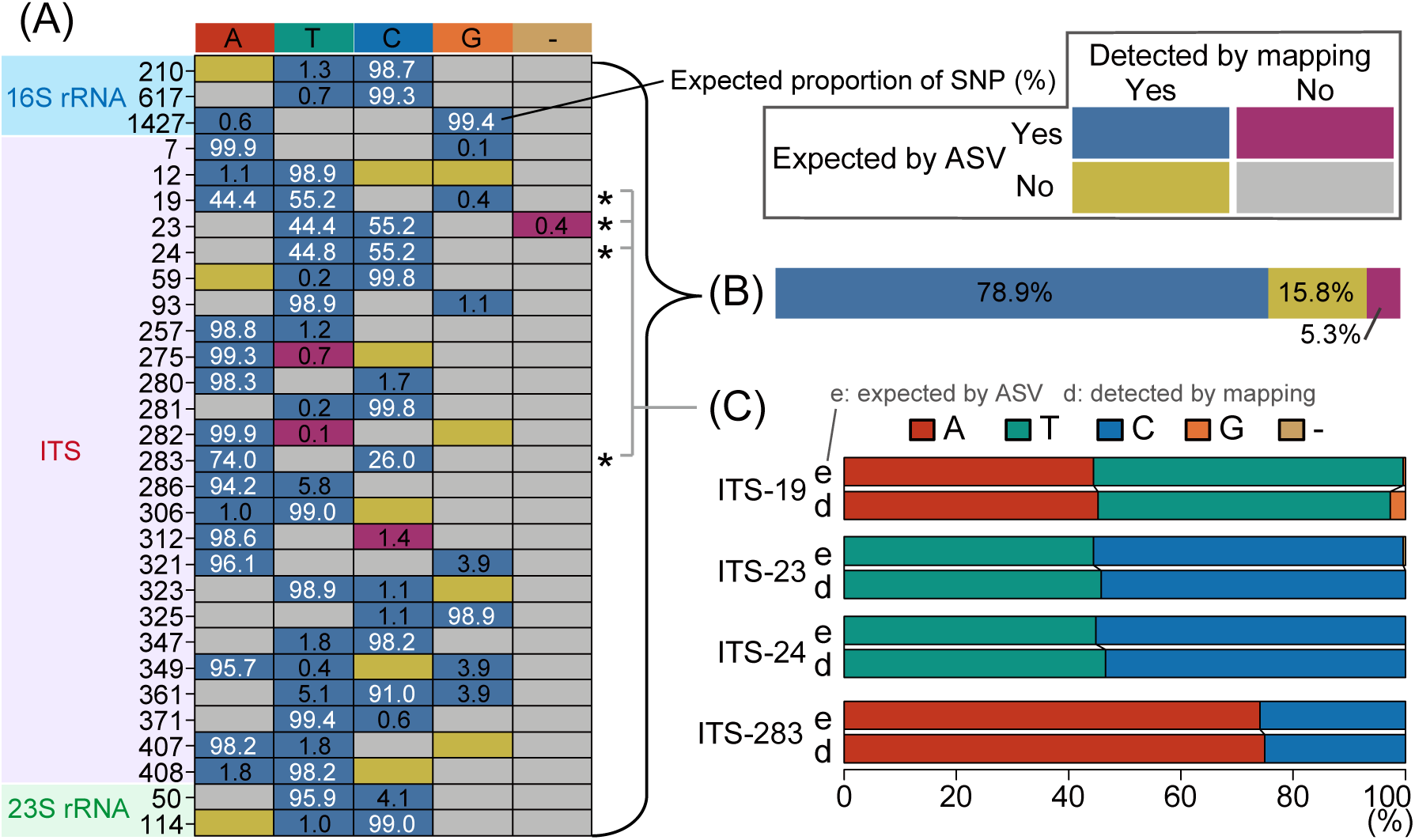
Comparison of single-nucleotide polymorphisms (SNPs) expected based on amplicon sequence variants (ASVs) and those detected through metagenomic read mapping of CL500-11 sequences from Lake Biwa. (A) The rows represent 30 individual SNP sites expected from ASVs in the lake. Row labels indicate base positions within the 16S rRNA, ITS, and 23S rRNA genes. Colors of the boxes indicate whether variations were expected from ASV, detected through mapping, or both (see the legend at the top right). Numbers in the boxes indicate the proportions (%) of SNPs expected from ASV abundances, where rare (<10%) variants are shown in black and others are in white. (B) The summarized proportions of expected and detected statuses among the observed SNPs illustrated in (A). (C) Comparison of the base proportions expected from ASV and detected through mapping for four major SNP sites (indicated by asterisks in (A)) that differentiate the four dominant ASVs in the lake (shown in Fig. 4).

Relatively low read throughput is another major challenge facing long-read amplicon sequencing. This limitation occurs because a significant number of reads are discarded during read quality control along with the lower per-cost sequencing throughput compared to short-read sequencing. In the present study, we used the stringent quality filtration threshold proposed for the original DADA2 pipeline developed for CCS-based long-read amplicon sequencing [29]. As a result, 28,153 reads were assigned to the ASVs, representing only 17.6% of the original CCS (generated using the ‘Minimum Predicted Accuracy=90’ parameter) and ∼11.3% of the nominal output of the sequencer (∼250,000 reads per five SMRT cells on PacBio RSII). Notably, another study targeting a shorter region (the full-length 16S rRNA gene) using a newer generation of sequencing platform (PacBio Sequel) and chemistry (S/P3-C3/5.0) reported more efficient read recovery using the same analytical pipeline [29]; that analysis (referred to as ‘Replicate 2’ of the fecal sample) resulted in 186,124 reads assigned to ASVs, representing ∼37.2% of the nominal output of the sequencer (∼500,000 reads per single SMRT cell on PacBio Sequel). The rapid improvement of sequencing and bioinformatics technologies will eventually overcome the limitation of throughput, making long-read amplicon sequencing one of the most promising approaches for high-resolution phylogenetic profiling of environmental microbial assemblages, as it complements the limited resolution of short-read amplicon sequencing and the limited sensitivity of the MAG-based approach.

The overall community composition in our study showed severe discrepancies compared to that described in a previous study [38] based on short-read amplicon sequencing of the same DNA extracts (Fig. S4). Whereas members of the phyla Actinobacteria and Chloroflexi were overrepresented in our long-read analysis, those affiliated with Bacteroidetes, Planctomycetes, and Verrucomicrobia were underrepresented (Fig. S4). Primer specificity could be one reason behind this effect. Although the primer 23Sr-mod shows improved universality over the original 23Sr, it still misses members of some bacterial groups and most Archaea (Table S2). For example, 23Sr-mod targets only 40.2% and 47.4% of all Planctomycetes and Verrucomicrobia, respectively, which might explain their underrepresentation in our results (Fig. S4). Another major factor affecting our results may be connectivity of the rRNA gene operon. A recent study reported that >40% of rRNA genes in environmental samples could be unlinked, meaning that the 16S and 23S rRNA genes are not in adjacent locations within a genome and thus the ITS region is absent [79]. Planctomycetes is one phylum with a large proportion of unlinked rRNA genes [79], which may have resulted in their underrepresentation (Fig. S4). While our approach can efficiently resolve microdiversity within individual lineages, it is not designed to reveal overall microbial community composition, for which conventional short-read amplicon sequencing or a direct quantification via fluorescence *in situ* hybridization is still a reasonable approach [80].

Finally, we note that genomic variation can exist even among cells with identical ITS sequences [15, 81, 82]. For example, the dominance of ASV_7 (the LD12 lineage) in both Lake Zurich and Japanese lakes (Fig. S1 and Supplementary Dataset) may have resulted from a shared ITS sequence among different genotypes, given the overall pattern of genetic disconnection between Japan and Europe (Fig. 3). Similarly, the unexpectedly low degree of diversification among CL500-11 sequences in Japanese lakes (Figs. 5 and S2) might be attributable to the inability to detect their genomic diversity using ITS sequences. Our method can detect differences in genotypes but cannot conclusively show homogeneity among them, and the latter characteristic remains to be tested in future works using genome-resolved approaches.

## Conclusions

In the present study, we applied single nucleotide-resolved long-read amplicon sequencing analysis to a large collection of environmental samples obtained from 11 deep freshwater lakes in Japan and Europe. The results demonstrated sympatric, allopatric, and temporal microdiversity in lake bacterioplankton and revealed phylogeographic patterns that could not be observed based on short-read amplicon sequencing or the MAG-based approach. Our results consistently supported genetic isolation between lakes in Japan and Europe, in contrast to previous reports of genomes sharing ANI>95% in freshwater habitats thousands of kilometers apart [40, 43, 44, 46] as well as in distant marine [83, 84] systems. The rapid accumulation of sequence data obtained from all over the world will allow for this topic to be revisited to draw broader conclusions about the global-scale dispersal and diversification processes of ubiquitous freshwater bacterioplankton lineages. Meanwhile, understanding intra-lineage population diversity at the regional scale (up to hundreds of kilometers), where dispersal limitation appears to be relatively weak, requires consideration of a complex combination of factors including the local environment, functional diversity of genotypes, migration frequency, genetic drift, and the ecological strategies of the lineage. The present study highlights the potential of long-read amplicon sequencing as a strategy for tackling these challenges. With the rapid improvement of sequencing technology, phylogenetic resolution beyond that of the 16S rRNA gene will become essential to microbial ecology and will reshape our understanding of environmental microbial diversity and ubiquity.

## Declarations

### Availability of data and material

The raw CCS reads generated in the present study were deposited under accession number PRJDB9651. The R script used in the DADA2 analysis workflow is available in the Supplementary Text. Read abundance tables and representative nucleotide sequences for the ASVs and OTUs are available in Supplementary Dataset.

### Authors’ contributions

YO, HT, and SN designed the research. YO, SF, MS, CC, AT, AK, and HO performed field sampling. YO analyzed data and wrote the draft. All authors contributed to finish the manuscript.

## Acknowledgements

We are grateful to T. Koitabashi, Y Goda, T Denboh, Y Ito, M Sugiyama, and E Loher for their assistance in field sampling.

