## Supplementary Information for "Microdiversity and phylogeographic diversification of bacterioplankton in pelagic freshwater systems revealed through long-read amplicon sequencing"

**This PDF file includes:**

Figures S1 to S4
Supplementary Text (R script)
Captions for Tables S1 and S2
Captions for Supplementary Dataset

**Other supplementary materials for this manuscript include the following:**

Tables S1 and S2
Supplementary Dataset

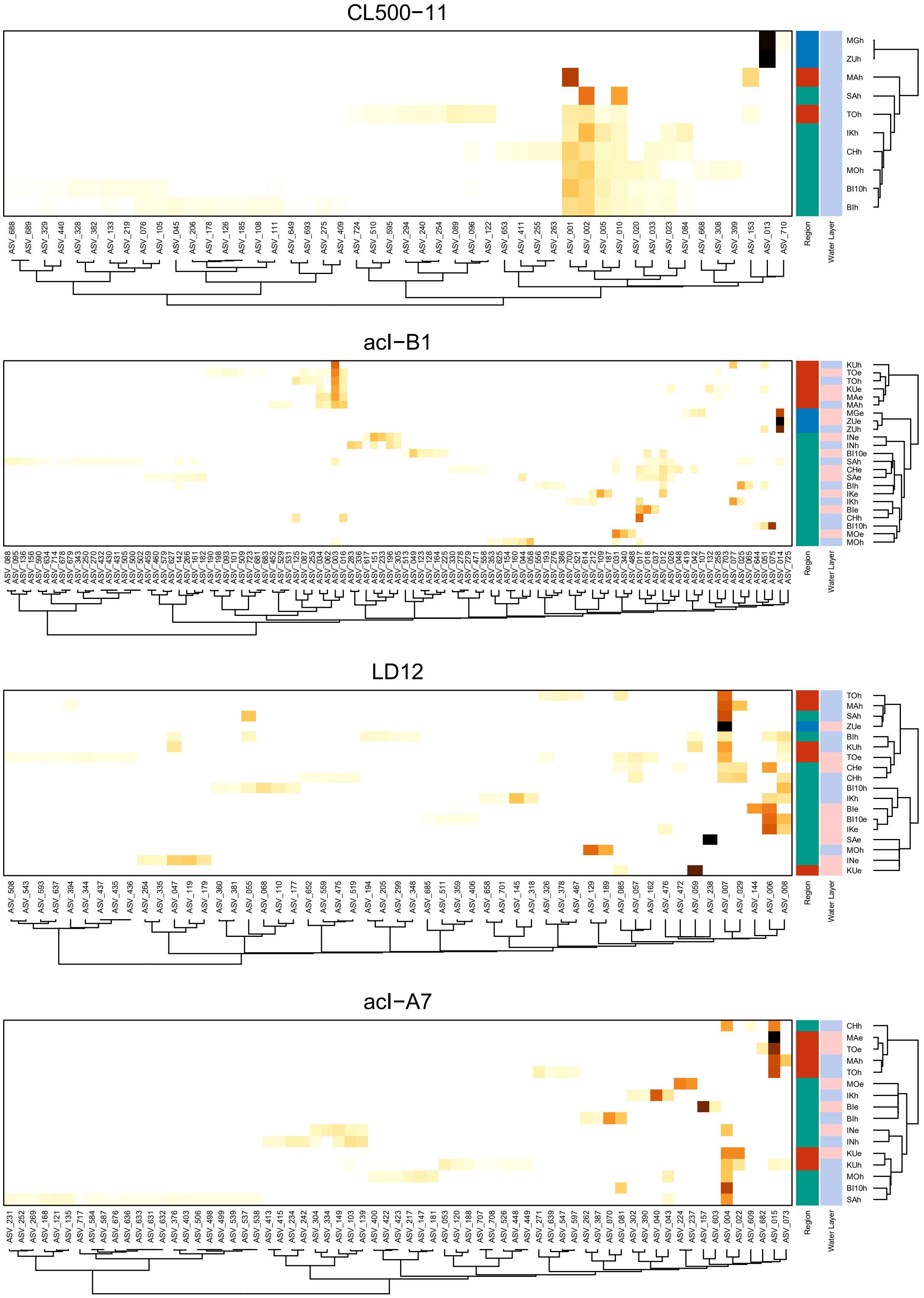


***Figure S1.*** The relative abundance of ASVs in each sample for each lineage. Rows and columns are clustered based on the Bray-Curtis dissimilarity among samples and ASVs, respectively (see Materials and Methods for detail). Abbreviations for sample names follow those in Fig. 3. Legends are shown at the bottom.

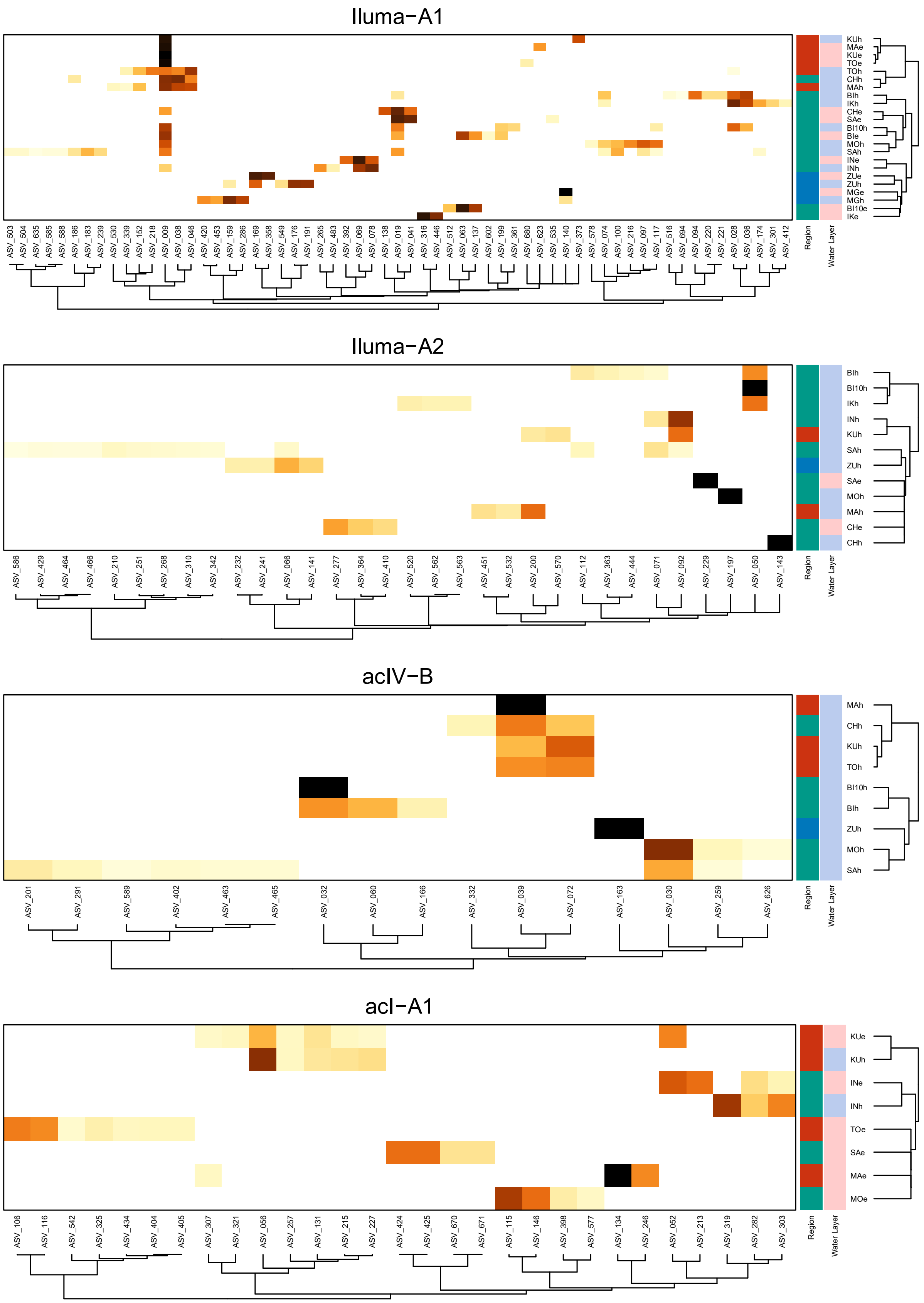


***Figure S1.*** continued.

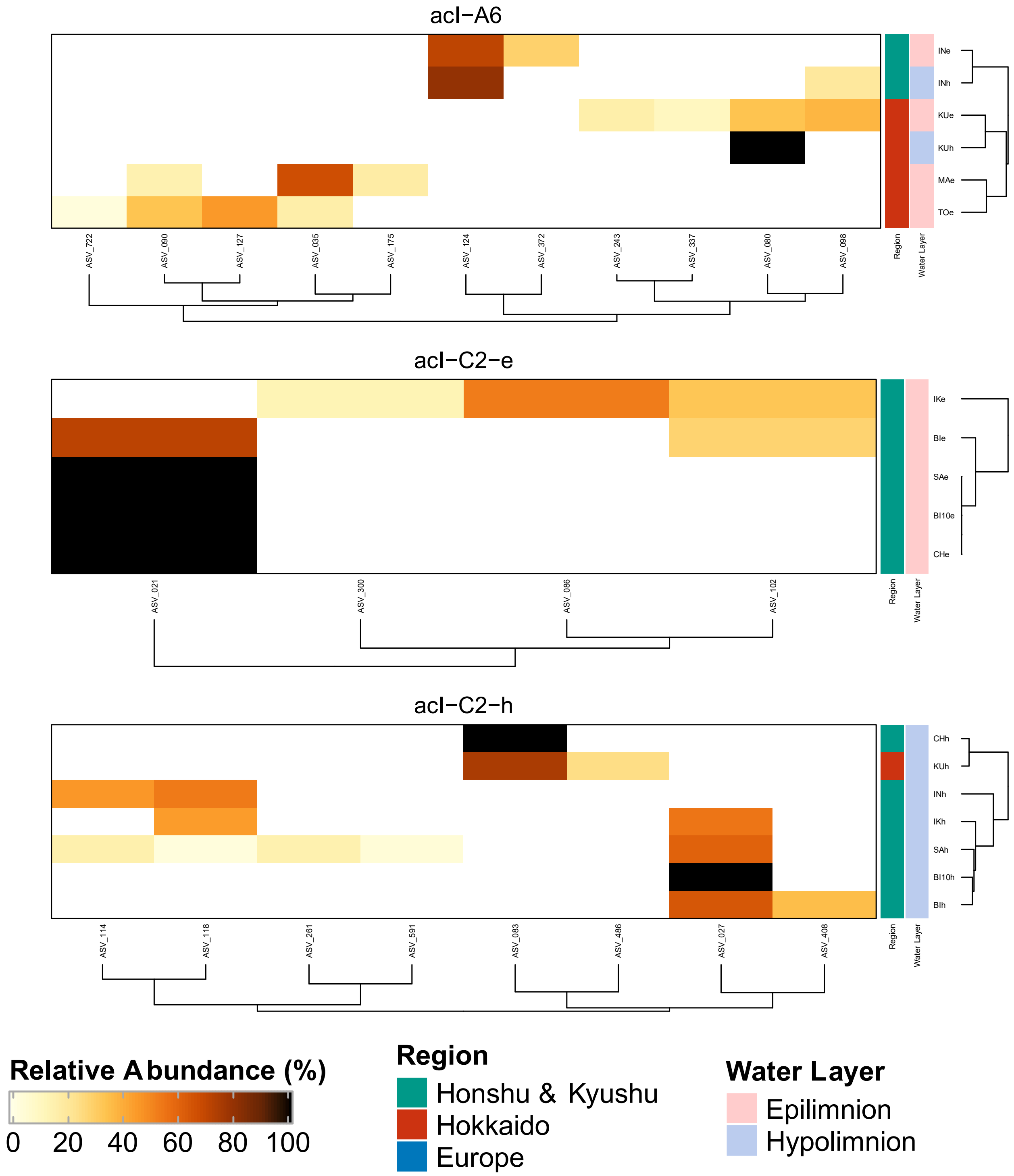


***Figure S1.*** continued.

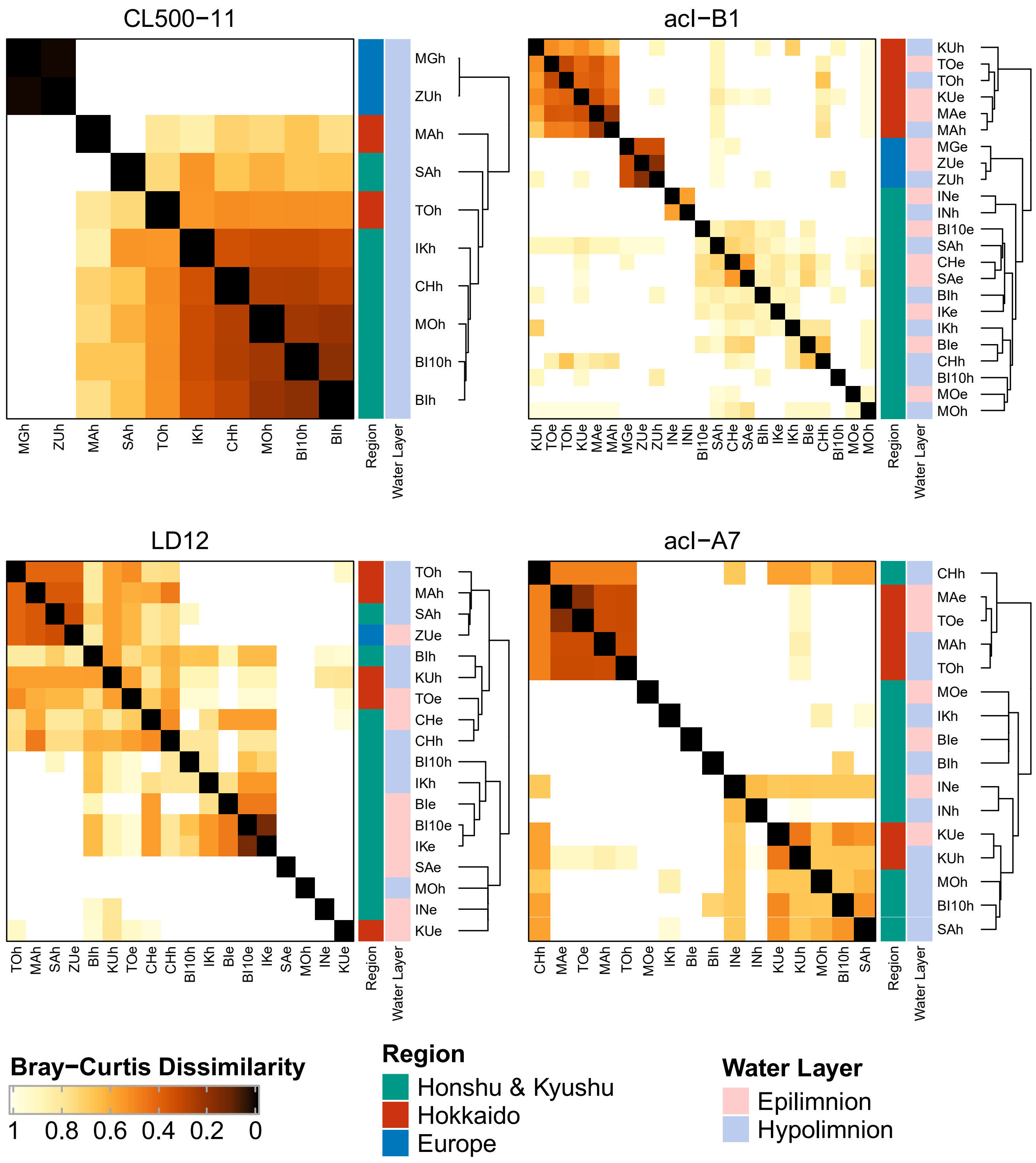


***Figure S2.*** Clustering of samples based on Bray–Curtis dissimilarity of amplicon sequence variant composition for each lineage. Abbreviations for sample names follow those in Fig. 3.

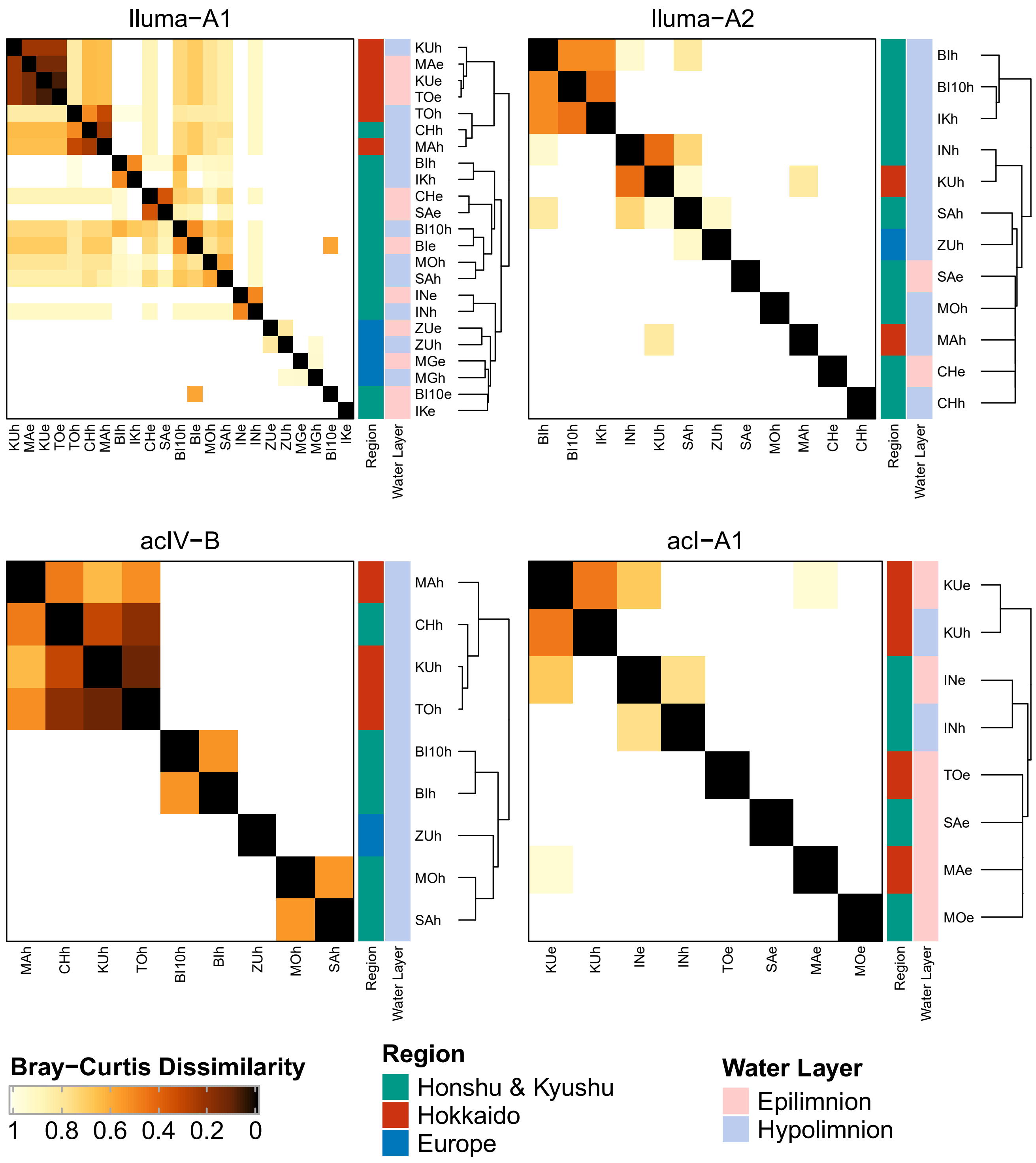


***Figure S2.*** Continued.

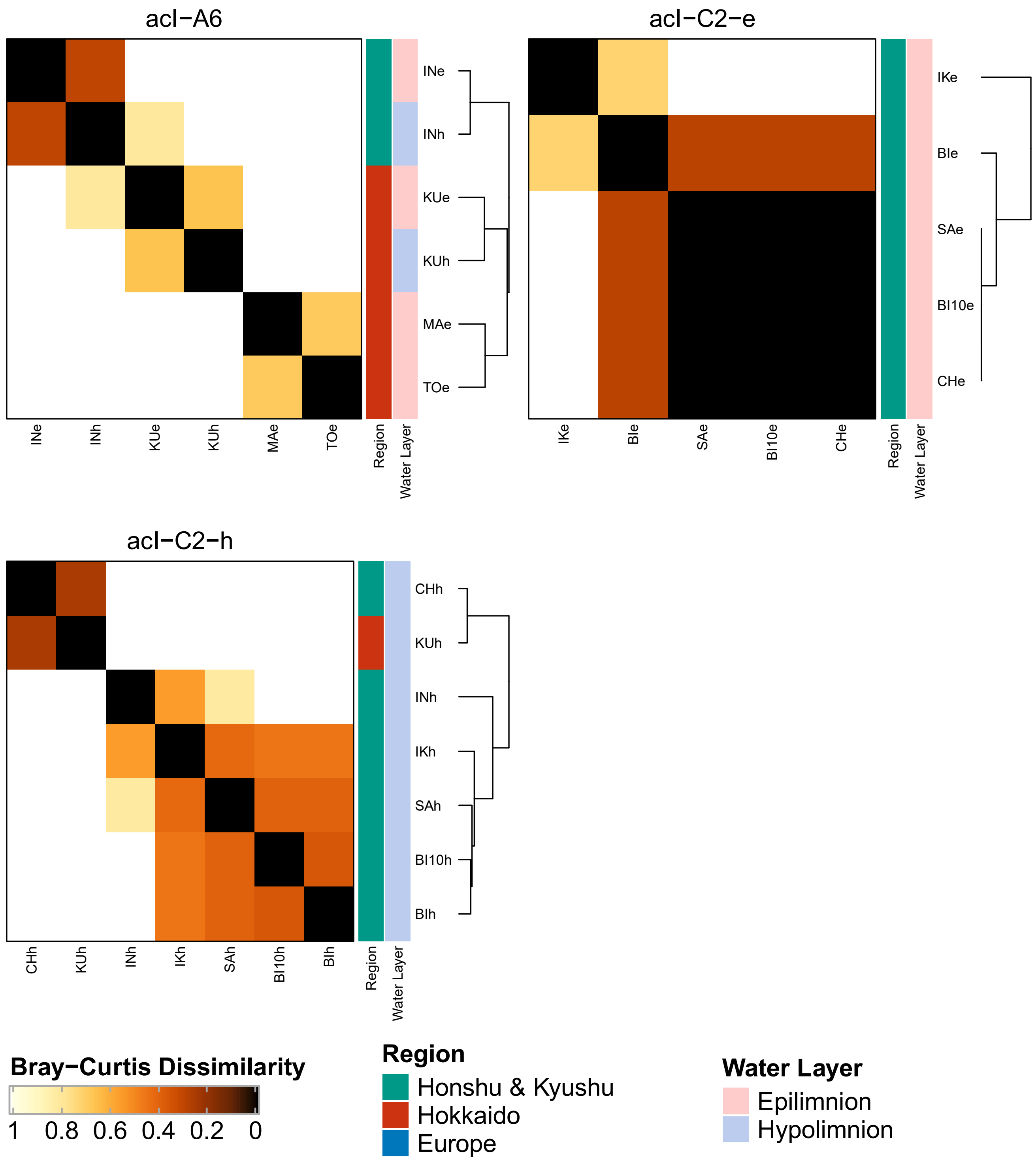


***Figure S2.*** Continued.

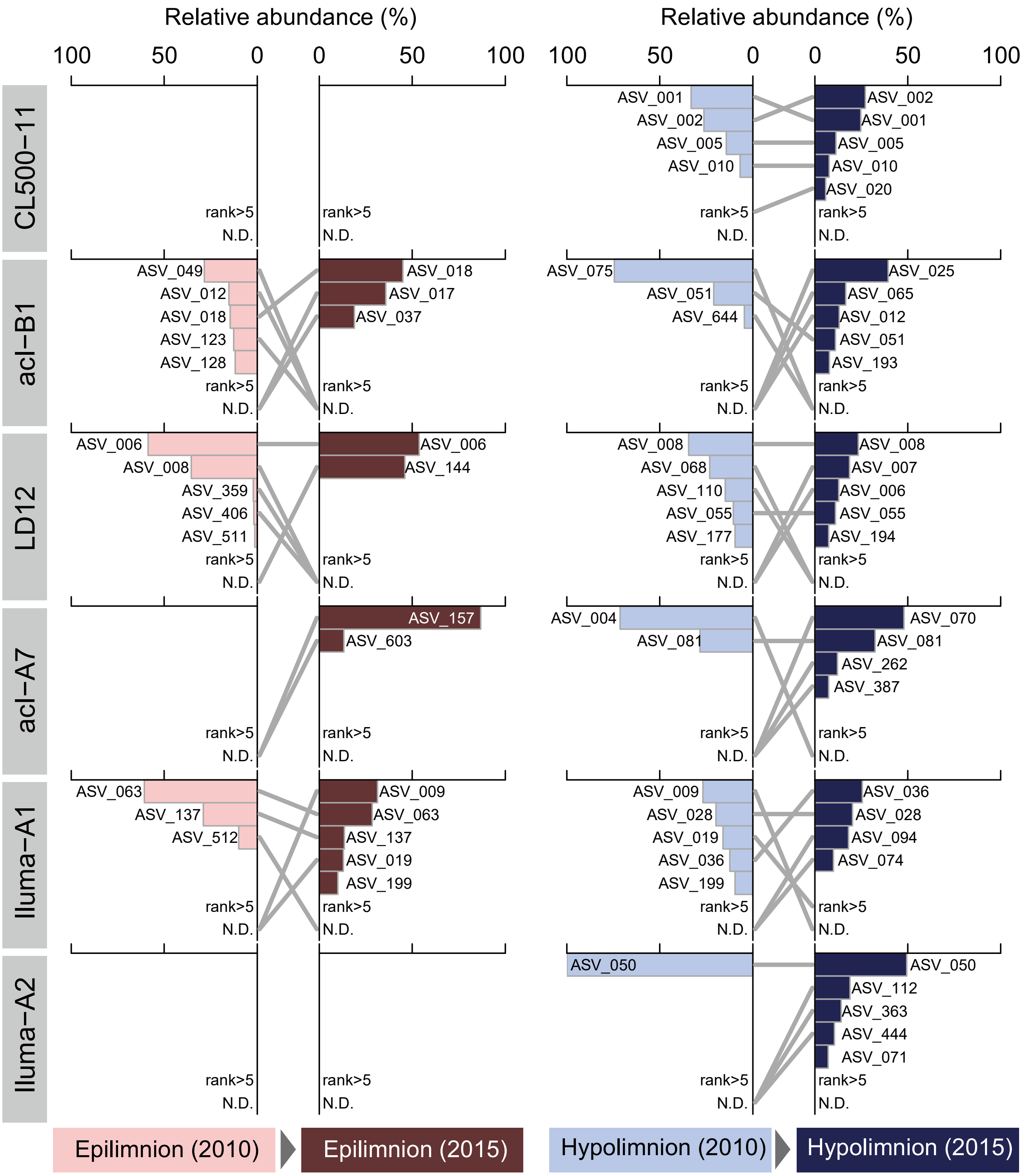


***Figure S3.*** Comparison of the five most abundant amplicon sequence variants (ASVs) between temporal replicates (2010 and 2015) collected in Lake Biwa for each water layer. Bars indicate the relative abundances of ASVs within each lineage and are ordered by abundance rank for each sample. Gray lines indicate succession of ranks between two time points; N.D., not detected.

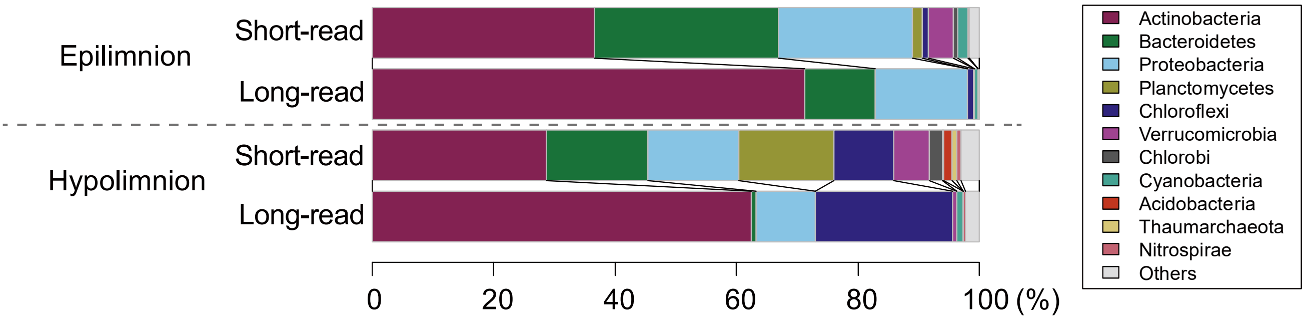


***Figure S4.*** Comparison of the taxonomic composition (at the phylum level) of reads generated using long-read (this study) and short-read (Okazaki et al., 2017) platforms. Data from nine Japanese lakes sampled in 2015 were averaged for both water layers. Note that the same DNA extracts were used for both studies.

### Supplementary Text (R script)

```
#The DADA2 pipeline used in the present study
##############################################
require("dada2")

#input raw fastq files
path<-"./00_rawfq"
fnFs<-sort(list.files(path,pattern=".fastq",full.names=TRUE))
sample.names<-sapply(strsplit(basename(fnFs),"[_,.]"),"[",2)

#input primer sequences
Fprimer<-"AGRGTTTGATYMTGGCTCAG"
Rprimer<-"RGTTBYCYCATTCRG"

#remove primers
nops<-file.path(".","01_noprimers",paste0(basename(fnFs),".gz"))
removePrimers(fnFs,nops,primer.fwd=Fprimer,primer.rev=dada2:::rc(Rprimer))

#filter reads
filtFs<-file.path("./02_filtered_reads",paste0(sample.names,"_hq.fq.gz"))
out<-filterAndTrim(nops,filtFs,maxEE=2,multithread=30,minQ=3,minLen=1500,maxLen=3000,rm.phix=FALSE)

#dereplication
drp<-derepFastq(filtFs,verbose=TRUE)

#learn error
err<-learnErrors(drp,errorEstimationFunction=PacBioErrfun,BAND_SIZE=32,multithread=30)

#save error
saveRDS(err, file.path(".", "err.rds"))

#denoise
dd2<-dada(drp,err=err,multithread=30)

#save dd2
saveRDS(dd2, file.path(".", "dd2.rds"))

#generate sequence table
st<-makeSequenceTable(dd2)

#de novo chimera removal
st.nochim<-removeBimeraDenovo(st,method="consensus",multithread=30,verbose=TRUE)
```

### Captions for other Supplementary Information Files:

**Supplementary Table S1**

Detailed profiles of the lakes sampled in the present study. Data were collected from Okazaki et al. (2017) and Okazaki et al. (2018).

† Sampling date and depths in 2010 are shown in parentheses.

**Supplementary Table S2**

Comparison of the 23Sr (original) and 23Sr-mod (modified for the present study) primers, showing their coverage for each phylum. Coverage was determined using the TestProbe 3.0 tool with reference to the SILVA LSU 132 Parc database (Quast et al., 2013) allowing no mismatches.

**Supplementary Dataset**

Summarized dataset related to amplicon sequence variants (ASVs) and operational taxonomic units (OTUs). The Excel file consists of two worksheets. In the first sheet (named "by_ASV"), each row represents an individual ASV. The columns indicate the corresponding SSU-ASV and OTU IDs, the taxonomy assigned to the OTU, and read abundance in each sample. The nucleotide sequences of the ASVs are shown in the last column. In the second sheet (named "by_OTU"), each row represents an individual OTU. The columns indicate the taxonomy, classification based on freshwater bacterioplankton nomenclature (see Methods for details), the number of SSU-ASVs and ASVs assigned to the OTU, the number of samples in which the OTU was detected, and read abundances in total and in each sample. OTUs are sorted based on the total read number. Representative 16S rRNA gene sequences of the OTUs are shown in the last column. Abbreviations for sample names follow those in Fig. 3.
